# Sub-chronic and acute toxicity of 6PPD-quinone to early-life stage rainbow trout (*Oncorhynchus mykiss*)

**DOI:** 10.1101/2024.09.25.614982

**Authors:** Catherine Roberts, Evan Kohlman, Niteesh Jain, Mawuli Amekor, Alper James Alcaraz, Natacha Hogan, Markus Brinkmann, Markus Hecker

## Abstract

N-(1,3-Dimethylbutyl)-N’-phenyl-p-phenylenediamine-quinone (6PPD-Q) is a derivative of rubber tires which leaches into surface waters when tire particles are swept into roadway runoff. 6PPD-Q has been identified as a potential cause of urban runoff mortality syndrome in coho salmon, and subsequent research has determined a wide species variation in toxicity among fishes. While adult rainbow trout are known to be sensitive, there is limited research on their early-life stages. Given that early-life stages of fish are often more sensitive than adults, the aim of these studies was to assess the acute and sub-chronic toxicity of 6PPD-Q in early-life stage rainbow trout. Rainbow trout alevins were exposed from hatch until 28 days post-hatch (dph) to time-weighted average 6PPD-Q concentrations ranging from 0.06-2.35 *μ*g/L. From these studies, a 28-day median lethal dose (LC_50_) of 0.56 *μ*g/L was derived. Morphological deformities were observed during the exposure period, including pooling of blood in the caudal fin. A follow-up acute study with exogenously feeding rainbow trout fry revealed a 96-hour LC_50_ of 0.47 *μ*g/L. These studies indicate that early-life stage rainbow trout are more sensitive than previously studied sub-adults, with exogenously feeding fry being the most sensitive life stage studied so far, and that sub-chronic exposure to 6PPD-Q can result in developmental abnormalities. This research highlights the importance of utilizing early-life stage studies to determine the most sensitive benchmark concentrations and their value in determining sub-lethal effects.

## INTRODUCTION

N-(1,3-Dimethylbutyl)-N’-phenyl-p-phenylenediamine-quinone (6PPD-Q) has become a chemical of very high concern since its identification as the likely causative agent of Urban Runoff Mortality Syndrome (URMS) in 2021. URMS, referring to the phenomenon of pre-spawn mortality incidents of coho salmon in the Pacific Northwest, has been linked to both precipitation events and proximity to roadways. Following years of investigation, Tian et al.(2021) found that 6PPD-Q, a transformation product of the tire antiozonant N-(1,3-Dimethylbutyl)-N’-phenyl-p-phenylenediamine (6PPD), is acutely toxic to coho salmon, with a 24-hr median lethal dose (LC_50_) of 0.095 *μ*g/L (Tian et al., 2022). The abrasion and subsequent deposition of tire wear particles (TWP) on road surfaces leads to pulsed emission of 6PPD-Q into aquatic ecosystems during rainfall events (Johannessen et al., 2021). Subsequent studies have reported environmental levels of 0.08-2.85 *μ*g/L of 6PPD-Q in urban water systems (Challis et al., 2021; Hu et al., 2022; Johannessen et al., 2021). These concentrations frequently surpass the toxicity threshold for coho salmon, as well as other salmonid species that have been discovered to be sensitive, including brook trout (*Salvelinus fontinalis*) with a 24-hr LC_50_ of 0.59 *μ*g/L (Brinkmann et al., 2022), white-spotted char (*Salvelinus leucomaenis pluvius*) with a 24-hr LC_50_ of 0.51 *μ*g/L (Hiki and Yamamoto, 2022), juvenile lake trout with a 96-hr LC_50_ of 0.50 *μ*g/L (Roberts et al., 2024), and sub-adult rainbow trout with a 96-hr LC_50_ of 1.00 *μ*g/L (Brinkmann et al., 2022). Many other species tested to date, covering other members of the salmonid family, as well as cyprinids and acipenserids, were found to be insensitive to 6PPD-Q exposure at environmentally relevant exposure concentrations (Brinkmann et al., 2022; Varshney et al., 2022).

The considerable variability in sensitivity among fishes to this compound is remarkable. While some salmonid species, such as *Salvelinus fontinalis*, exhibit sensitivity at low concentrations of 6PPD-Q, other closely related species of the same genus, including Arctic char (*Salvelinus alpinus*) and southern Asian dolly varden (*Salvelinus curilus*), show no acute adverse effects even at very high concentrations (Brinkmann et al., 2022; Hiki and Yamamoto, 2022). Nevertheless, there is a lack of experimental data regarding the acute lethality following exposure to 6PPD-Q for most salmonid (and non-salmonid) species, and currently, there are no reliable indicators enabling risk assessors to predict or otherwise identify species as vulnerable to this chemical.

Moreover, there is limited knowledge regarding the early-life stage or chronic exposure of salmonid species to 6PPD-Q. Given that embryonic and larval life stages of fish are often more sensitive to contaminants than adults of the same species, these life stages may be more susceptible to lethal or sub-lethal effects than their adult counterparts. The results of acute adult exposures may not be protective of earlier life stages nor indicate sub-lethal effects during growth and development (Hutchinson et al., 1998) – an oversight that can have significant ramifications for wild populations of fish, as early-life-stage mortality can threaten future generations. Considering that 6PPD-Q can be expected to continue to leach from TWPs once these particles are deposited in sediments, there is a potential risk that salmonid alevins, which inhabit gravel for the first stages of life, may be particularly susceptible to exposure to 6PPD-Q. While aqueous exposure to 6PPD-Q through urban stormwater runoff typically occurs in pulses, it is possible that exposure of salmonid alevins to TWPs in sediments could happen over extended exposure periods.

This study aimed to determine both sub-chronic and acute toxicity of 6PPD-Q to early-life stage rainbow trout. Rainbow trout are an important model species in the aquatic hazard assessment of chemicals (OECD, 2000; Teather and Parrott, 2006; Thorgaard et al., 2002), and subadult individuals of this species were previously found to be sensitive to 6PPD-Q exposure (Brinkmann et al., 2022). Here, we studied the sub-chronic toxicity of 6PPD-Q to rainbow trout alevins, beginning at hatch and continuing until the transition to the exogenously feeding fry stage is completed. We also studied the lethality of 6PPD-Q to exogenously feeding fry during a 96-hr acute exposure. In addition to mortality, sub-lethal effects on growth, development, and histopathology were studied that might have implications for fish health and population dynamics. This study provides critically needed information on 6PPD-Q toxicity to environmental risk assessors and chemical managers.

## MATERIALS AND METHODS

### Chemical source

Neat 6PPD-Q was sourced from Toronto Research Chemicals (Cat# P348790, Lot # 6-ABK-93-1, purity 97%). Working stocks were prepared using dimethyl sulfoxide (DMSO) as the solvent, at 10,000× the final exposure concentrations, to achieve a final concentration of 0.01% DMSO (v/v) in all treatments. A mass-labelled internal standard (6PPD-Q d5) (Cat#P348691) was also obtained from Toronto Research Chemicals and dissolved in LC-MS-grade methanol at a concentration of 1 mg/L. Fresh exposure solutions were prepared daily.

### Rainbow trout source and housing

Eyed rainbow trout embryos of triploid females were obtained from Lyndon Hatcheries (ON) and maintained in glass aquaria until hatch. Aquaria were kept at 14°C, with a 16:8 light:dark schedule, aerated, with a pump for water movement. All experiments involving the use of animals were reviewed and approved by the University of Saskatchewan Animal Research Ethics Board under Animal Use Protocol 20220002.

### Sub-chronic exposure

The concentrations of 6PPD-Q for the early-life stage study were determined based on the results of a previous acute exposure experiment with subadult rainbow trout, which showed the 96-hr LC_50_ in that life stage to be 1.00 *μ*g/L (Brinkmann et al. 2022). Nominal concentrations used were 4, 2, 1, 0.5, 0.25, and 0.125 *μ*g/L, along with a control group consisting of 0.01% DMSO in water. Each tank (2.5-L acrylic tanks) represented one replicate and was filled with exposure water for a 24-hr acclimation period prior to initiation of exposure. Each tank was individually aerated and placed randomly on shelving. All treatment groups were replicated five times. Tanks were maintained at a water temperature of 14 ± 0.5°C, and under a light:dark schedule of 16:8. The exposure began upon hatch, with 18 alevins randomly allocated to each replicate. Following 96 hours of exposure, five alevins per tank were sub-sampled for later transcriptomics analysis as a part of a parallel study. The exposure ran for 28 days, during which daily checks for mortalities and deformities were carried out during a 70% water change of each tank. As the alevins reached swim-up, they were fed brine shrimp (*Artemia* nauplii) once per day, and the tanks were culled to a maximum of ten fry each. At one week post-swim up, feeding was increased to twice daily for the remainder of the experiment, save for a 24-hr fasting period prior to the final sampling on day 28. Weekly water quality measurements of dissolved oxygen, ammonia, nitrate, nitrite, pH, and hardness were taken. At the end of the experiment, fry were euthanized using 150 mg/L buffered tricaine mesylate (MS-222), weighed, and measured for total and standard length.

### Juvenile acute exposure

After the conclusion of the sub-chronic exposure, rainbow trout fry from the same batch of embryos which were unexposed, now six weeks post-hatch (6wph), were exposed to 6PPD-Q for 96 hours. Nominal concentrations of 0.5, 1, and 4 *μ*g/L of 6PPD-Q plus a 0.01% DMSO control were used, employing the same experimental design as described above. Fry were exposed at the same temperature and lighting schedule as those in the sub-chronic study and were not fed over the exposure period. Fry were euthanized in 150 mg/L of buffered MS-222, weighed, and total and standard lengths taken.

### Deformity Analysis

Alevins were observed daily throughout the exposure for any morphological deformities, including yolk sac edema, spinal curvature, and craniofacial defects as described in Holm et al. (2003). Pooling of blood was also observed in the caudal fin and eye, and subsequently included in daily checks. Deformities were examined under the dissection microscope (ZEISS Stemi 208), photographed (ZEISS Axiocam 105 color, Labscope), and recorded. No severity scoring was performed, and occurrence was not normalized to mortality.

### Histopathology protocol

Samples for histopathology were fixed in 10% buffered formalin for 48 hours, and then stored in 70% ethanol. Samples were sent to Prairie Diagnostic Services (University of Saskatchewan, Saskatoon, Canada) for processing, sectioning, and staining. Fish were trimmed behind the operculum to separate the head from the body, dehydrated, and fixed in paraffin wax. Samples for gut analysis were sectioned thrice along the transverse plane, 200 *μ*m apart and stained with hematoxylin and eosin. Samples for gill analysis were sectioned along the sagittal plane, with three sections taken at 40*μ*m intervals and stained with hematoxylin and eosin. One fish from each replicate was sectioned for gut and gill samples of solvent control, 0.5 *μ*g/L, and the 1 *μ*g/L treatments. Because fewer fish survived the exposure in the 2 *μ*g/L concentration, five fish total were sampled across three reps for this treatment. Sectioning for both gills and gut did not provide consistent framing of organs, and as such, only qualitative assessments were performed.

### Chemical analysis

Samples of exposure water were collected at a time 0 hr (following a water change) and time 24 hr (just prior to a water change), at either three (acute study) or four (chronic study) separate time points throughout the experiment, to quantify 6PPD-Q losses. Water samples were frozen at – 20°C until analysis. For confirmation of concentration, 950 *μ*L of water was sampled from each tank, and 50 *μ*L of deuterium-labeled internal standard solution (1 mg/L) added. Analysis was performed using ultra-high-performance liquid chromatography and high-resolution mass spectrometry as previously described by Challis et al. (2021)The calibration standards used were within 15% of nominal concentrations, and TraceFinder 4.1 was used for target quantification. Concentrations across all replicates in a treatment were reported as a time-weighted average (TWA).

### Data analysis and statistics

The Kaplan-Meier function S_t+1_ = St*((N_t+1_-D_t+1_)/N_t+1_) was used to calculate percent mortality, and LC_50_s were interpolated using the [agonist] vs. response – variable slope (four parameters) model using Prism 10.1.2 for Windows (GraphPad Software, Boston, MA). Standard length and weight were analyzed using a nested one-way ANOVA using Prism 10.1.2 for Windows (GraphPad Software, Boston, MA).

## RESULTS AND DISCUSSION

### Exposure concentration validation and water quality parameters

Measured 6PPD-Q concentrations ranged from 39-64% of the nominal concentration, with an average 24-hour loss of 34.0%. This loss is slightly greater than that observed in other studies (Brinkmann et al., 2022; Greer et al., 2023). This increased loss may be due to a greater biomass-to-water ratio, thus increasing the biological capacity to metabolize the 6PPD-Q, or due to sorption to exposure materials, such as into tubing and tank walls. Time-weighted average concentrations of 6PPD-Q were 0.06, 0.10, 0.20, 0.44, 1.30, and 2.35 *μ*g/L for the sub-chronic exposure, and 0.26, 0.61, and 2.40 *μ*g/L for the 96-hr juvenile exposure. All results and analyses are reported based on the measured TWA concentrations.

Water quality parameters were as follows: 0.08 ± 0.09 mg/L ammonia, 0.21 ± 0.08 mg/L nitrite, 0.66 ± 0.25 mg/L nitrate, 60 ± 8.6 mg/L total hardness, 8.57 ± 0.50 pH, and 90.0 ± 4.6 % dissolved oxygen. These parameters are within the limits prescribed by international guidelines for juvenile rainbow trout, such as the Organization for Economic Cooperation and Development (OECD) test guideline No. 215 (OECD, 2000).

### Sub-chronic mortality

This study found a 28-day LC_50_ of 0.56 [95% confidence interval (CI) of 0.48 to 0.66 *μ*g/L] (Figure 2) for early-life stage rainbow trout. In comparison, the acute 24-hr LC_50_ for juveniles of coho salmon, the most sensitive species identified thus far, was 0.041 *μ*g/L (Lo et al., 2023), while a more comparable sub-chronic early-life stage study found a 45-day LC_50_ for lake trout juveniles of 0.39 *μ*g/L (Roberts et al., 2024). Rainbow trout early-life stages are therefore at similar risk as lake trout juveniles, and while this value is representative of a longer timeframe, 0.56 *μ*g/L is an environmentally relevant concentration of 6PPD-Q (Challis et al., 2021) and is a lower threshold for mortality than reported for the majority of the sensitive adult species studied thus far (Brinkmann et al., 2022; Hiki and Yamamoto, 2022). Significant mortality of rainbow trout began four days post-initiation of exposure and continued throughout the exposure (Figure 1). The delay in time to mortality is interesting, as in acute adult studies, mortality in sensitive species occurred within hours of exposure (Brinkmann et al., 2022; Tian et al., 2021). This is, however, consistent with findings of a study conducted with lake trout alevins, also beginning at hatch, where mortality was evident on a timescale of days rather than hours (Roberts et al., 2024). This may be due to changes in respiratory function, where newly hatched alevins largely respire through their skin until the gills are more developed (Rombough and Ure, 1991). This suggests that the gills are a key organ in toxicity. In treatments where significant mortalities were observed, these predominantly occurred within the first two weeks of exposure, with the highest treatment of 2.35 *μ*g/L exhibiting the steepest mortality between days 4 and 12. Interestingly, the remaining three fry in this treatment survived for the remainder of the exposure period. Similarly, the 0.44 *μ*g/L fry that did not succumb within the first two weeks survived to the completion of the exposure. No overt symptoms consistent with URMS (Tian et al., 2022) or previous laboratory studies with sub-adult fish (Brinkmann et al., 2022) were observed in this experiment, although no systemic assessment of behaviour was conducted.

**Figure 1.**
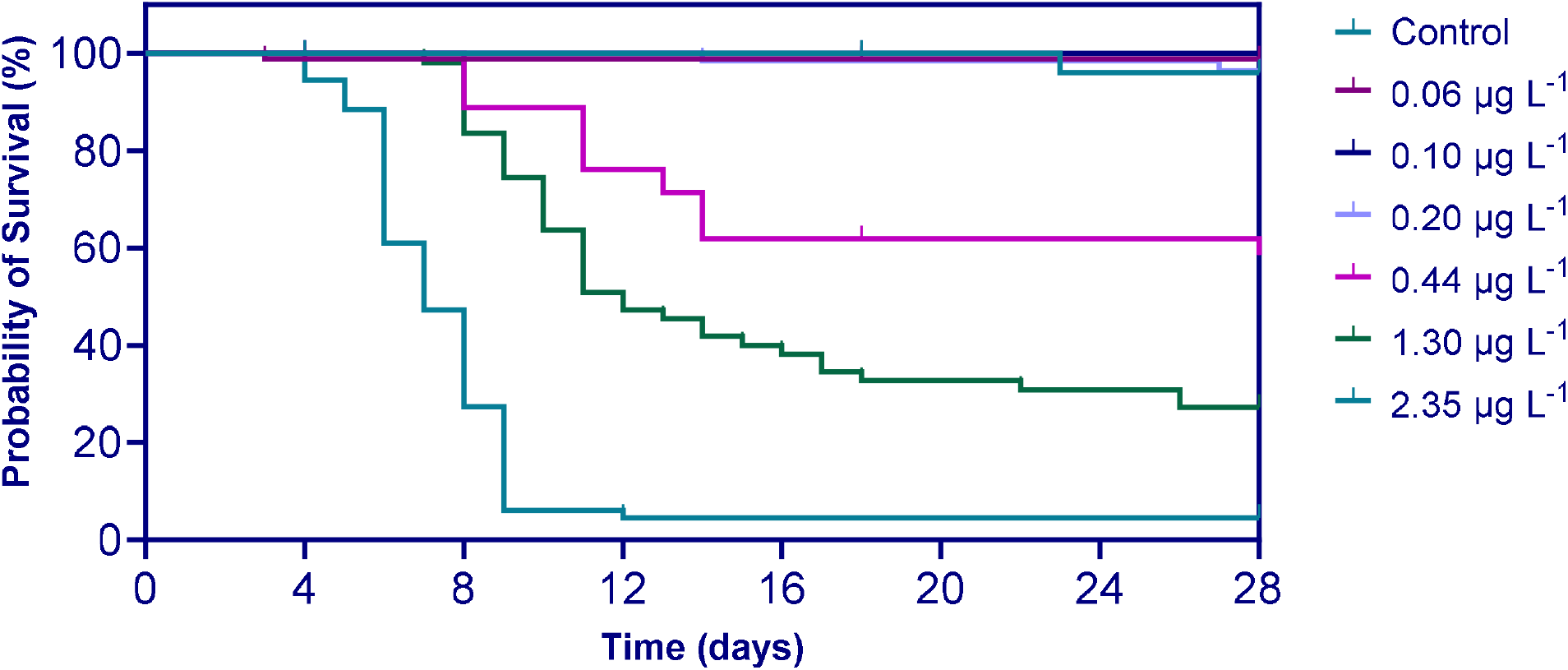
Survival time analysis for rainbow trout exposed to 6PPD-Q beginning at hatch and continuing for 28 days. Concentrations are time-weighted average measured concentrations.

**Figure 2.**
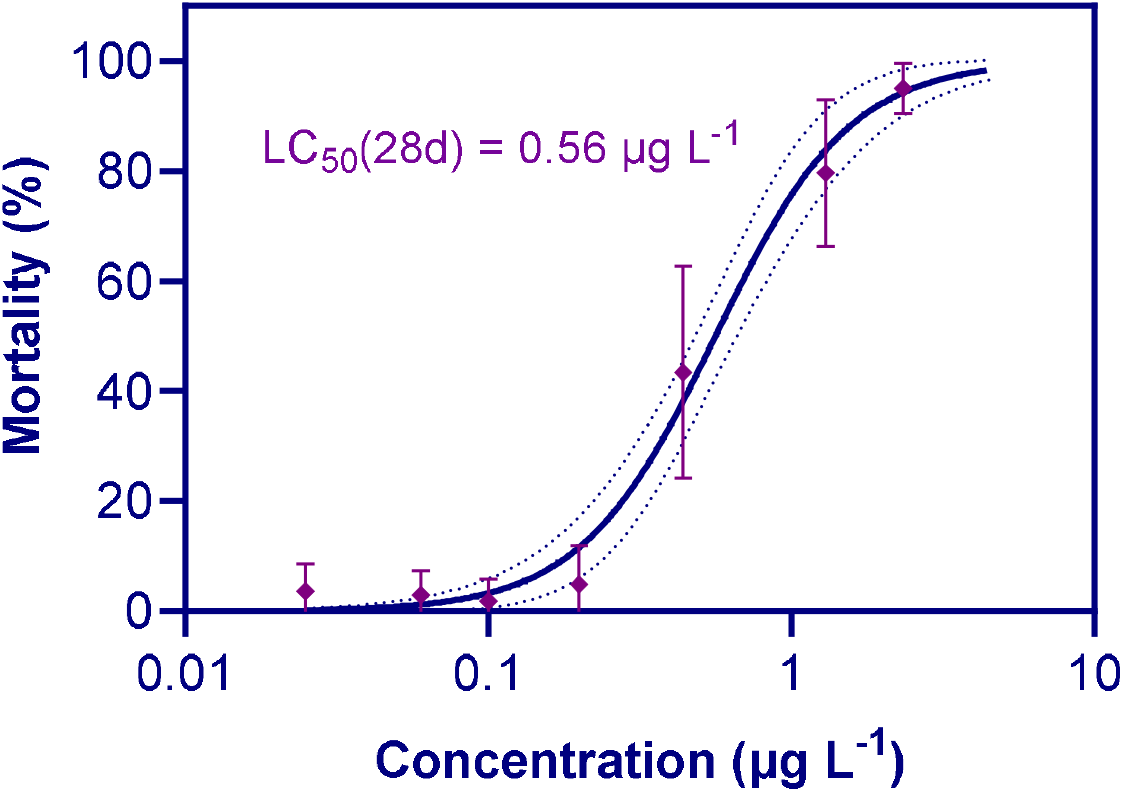
Percent mortality of rainbow trout exposed to 6PPD-Q beginning at hatch and continuing for 28 days, with a 28-day LC_50_ 95% confidence interval (CI) of 0.48 to 0.66 *μ*g/L. Points represent mean of replicates for each concentration, standard deviation is represented by the bars. The dotted lines indicate the 95% confidence interval. Concentrations on the x-axis are based on time-weighted average measured concentrations.

### Sub-chronic gross pathology

No significant difference in standard length of those fish which survived to the completion of the exposure was found among treatments, and only the 1.30 *μ*g/L treatment exhibited a significant difference in total length (S1, S2). Similarly, a significant difference in weight was found only in the 2.35 *μ*g/L treatment (S3). However, given the small surviving sample size of the highest two treatments upon termination of exposure, this is not necessarily attributed to exposure to 6PPD-Q. This treatment had only three individuals remaining at the termination of the exposure and, thus, did not have a representative sample size.

While acute and sub-chronic mortality was evident in these life stages, the sub-lethal effects were also significant. In the sub-chronic study, deformities occurred in several treatment groups, including yolk sac edema, spinal curvature, and pooling of blood into the caudal fin (Figure 3). Given that these effects were only observed in the sub-chronic study, these pathologies are likely related to the disruption of key processes during later embryonic/alevin growth and development. Changes in the skeletal structure of the caudal fin may play a role in the observed pooling of blood in the caudal fin. As described by Greer et al. (2023), 6PPD-Q exposure in coho embryos resulted in changes in the expression of genes which play a role in bone development. Disruption of these genes at a critical time of growth and development may have inhibited the proper development of the skeletal structure of these fry, altering the flow of blood through the caudal fin. The pooling of blood may also be linked to the changes in vascular permeability, which has been observed following 6PPD-Q exposure both *via* changes in gene expression in vascular pathways (Greer et al., 2023), as well as in changes to the blood-brain barrier in exposed coho (Blair et al., 2021). Previous *in vitro* work suggests mitochondrial disruption as a potential mechanism of action (Mahoney et al., 2022), which can result in the production of reactive oxygen species. These reactive oxygen species can impact the permeability of the vasculature, not only around the brain, as is purported to be a contributor to 6PPD-Q toxicity (Blair et al., 2021; Liao et al., 2024), but also of the caudal fin of these fry.

**Figure 3.**
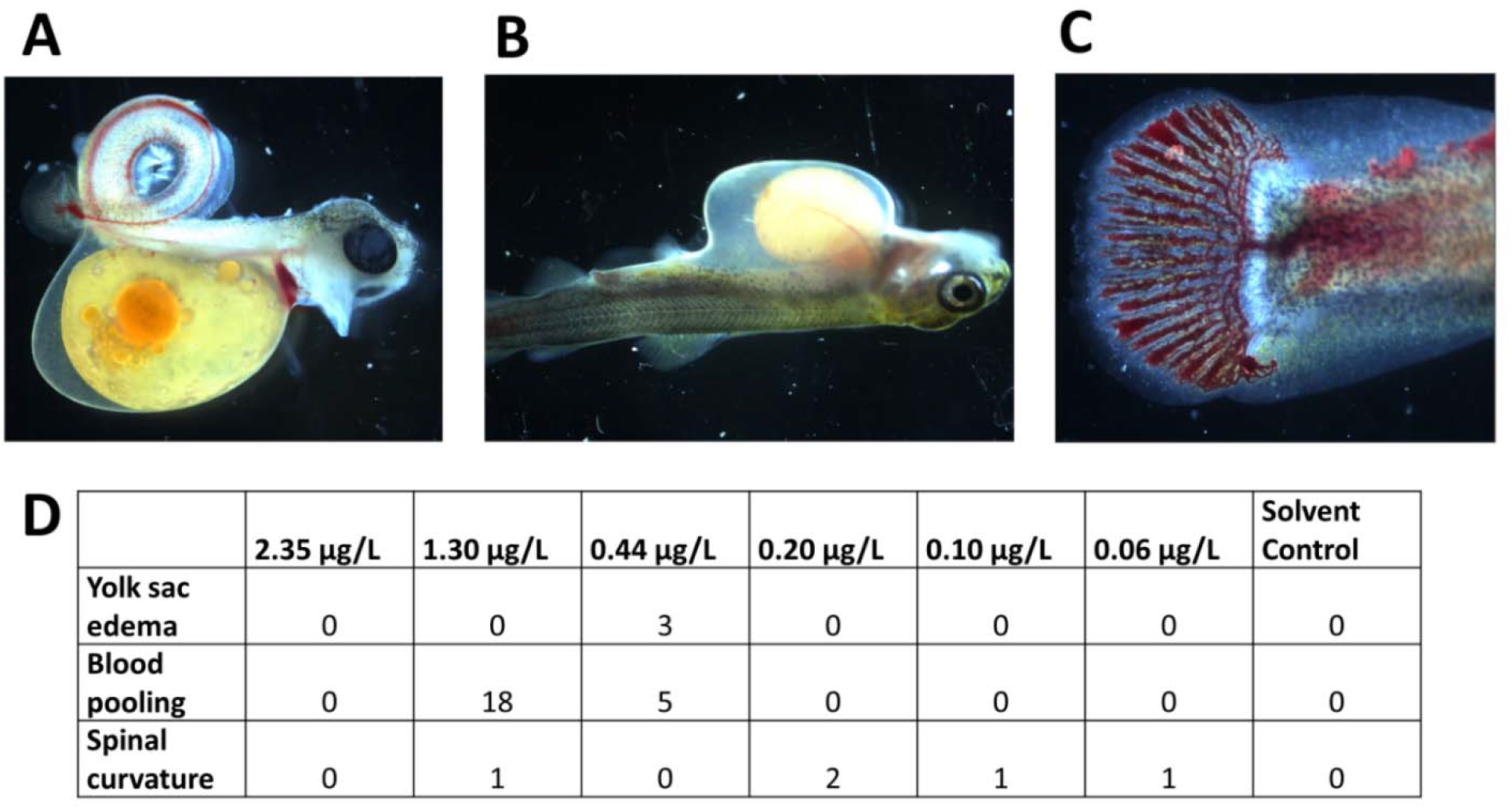
Examples of morphological changes in rainbow trout alevins exposed to 6PPD-Q from for 28 days from hatch. **A**. Spinal curvature occurring at 2.35 *μ*g/L **B**. Yolk sac edema occurring at 0.44 *μ*g/L **C**. Caudal fin pathology occurring at 0.44 *μ*g/L. **D**. Total incidences of morphological abnormalities over 28 days in each treatment (note: these represent total numbers and were not normalized to mortalities).

In addition to gross pathologies, histopathological assessment found qualitative changes in gill structure, including reduced length of gill filaments (Figure 4). Changes in the structure of the gill, as well as the delay in time to mortality observed in the sub-chronic exposure, indicate the gill as an important organ in toxicity. The behavioural changes observed during the acute exposure (gasping, surface swimming, loss of coordination) lend credence to this position, as these symptoms are also indicative of hypoxia. However, while many of these findings point toward this conclusion, more research is needed to determine the mechanism(s) at work, and what other sub-lethal other life stages and species may experience.

**Figure 4.**
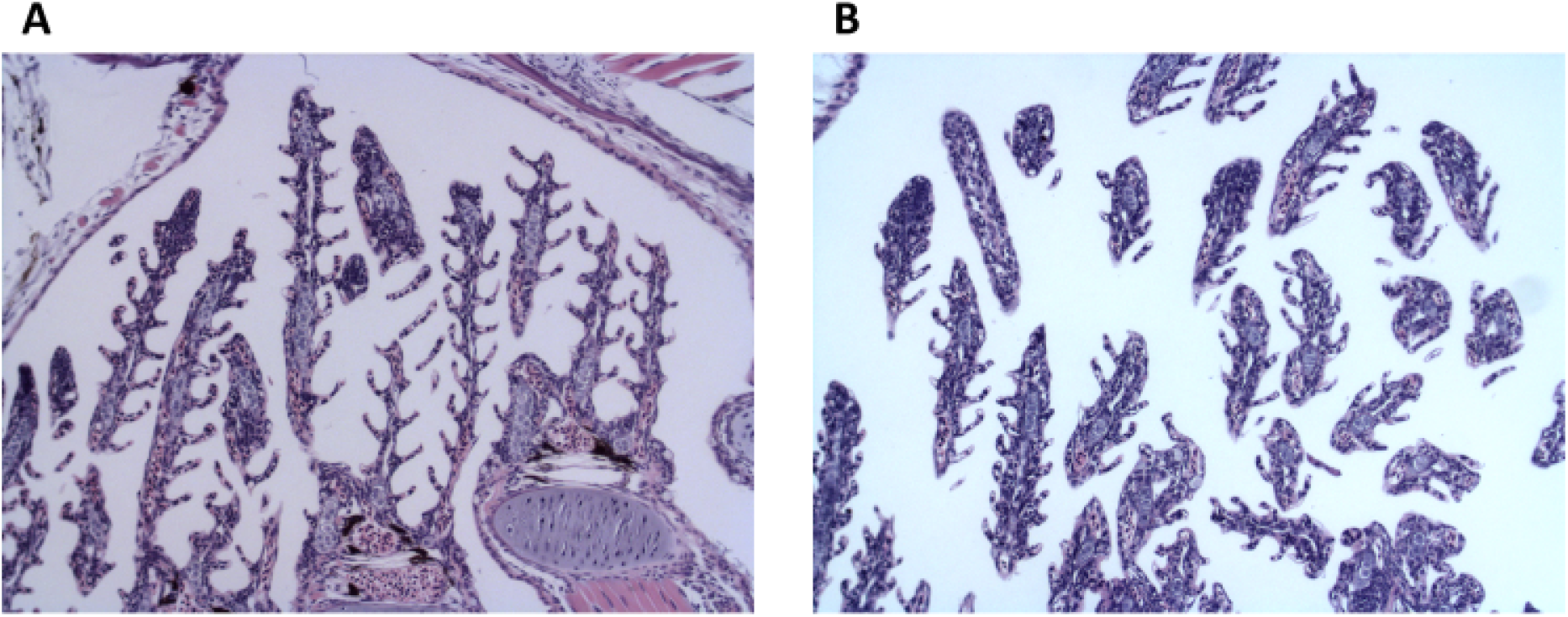
Example of histology of gills from rainbow trout alevins exposed to 6PPD-Q for 28 days from hatch, stained with hematoxylin and eosin, taken at 10× magnification. **A**. Solvent control (0.01% dimethyl sulfoxide). **B**. 1.30 *μ*g/L 6PPD-Q.

### Acute exposure

Exposure of exogenously feeding fry (six weeks post-hatch) to 6PPD-Q found significant and concentration-dependent mortalities at environmentally relevant concentrations (Figure 5). The 96-hr LC_50_ for fry was 0.47 *μ*g/L (95% CI of 0.42 to 0.60 *μ*g/L). This is in contrast with the study conducted on sub-adult rainbow trout, which found a 96-hr LC_50_ of 1.00 *μ*g/L (95% CI of 0.95 to 1.05 *μ*g/L) (Brinkmann et al., 2022). However, increased sensitivity in free-feeding fry versus sub-adults is consistent with findings in coho salmon (Lo et al., 2023; Tian et al., 2021). Changes in the behaviour of exposed fish were noted within four hours of initial exposure, with fry exhibiting gasping, loss of coordination, and surface swimming (S4). These behaviours are consistent with observations in coho salmon (Tian et al., 2021), rainbow trout, and brook trout sub-adults (Brinkmann et al., 2022), as well as exogenously feeding lake trout fry (Roberts et al. 2024). None of the fish exhibiting symptoms recovered.

**Figure 5.**
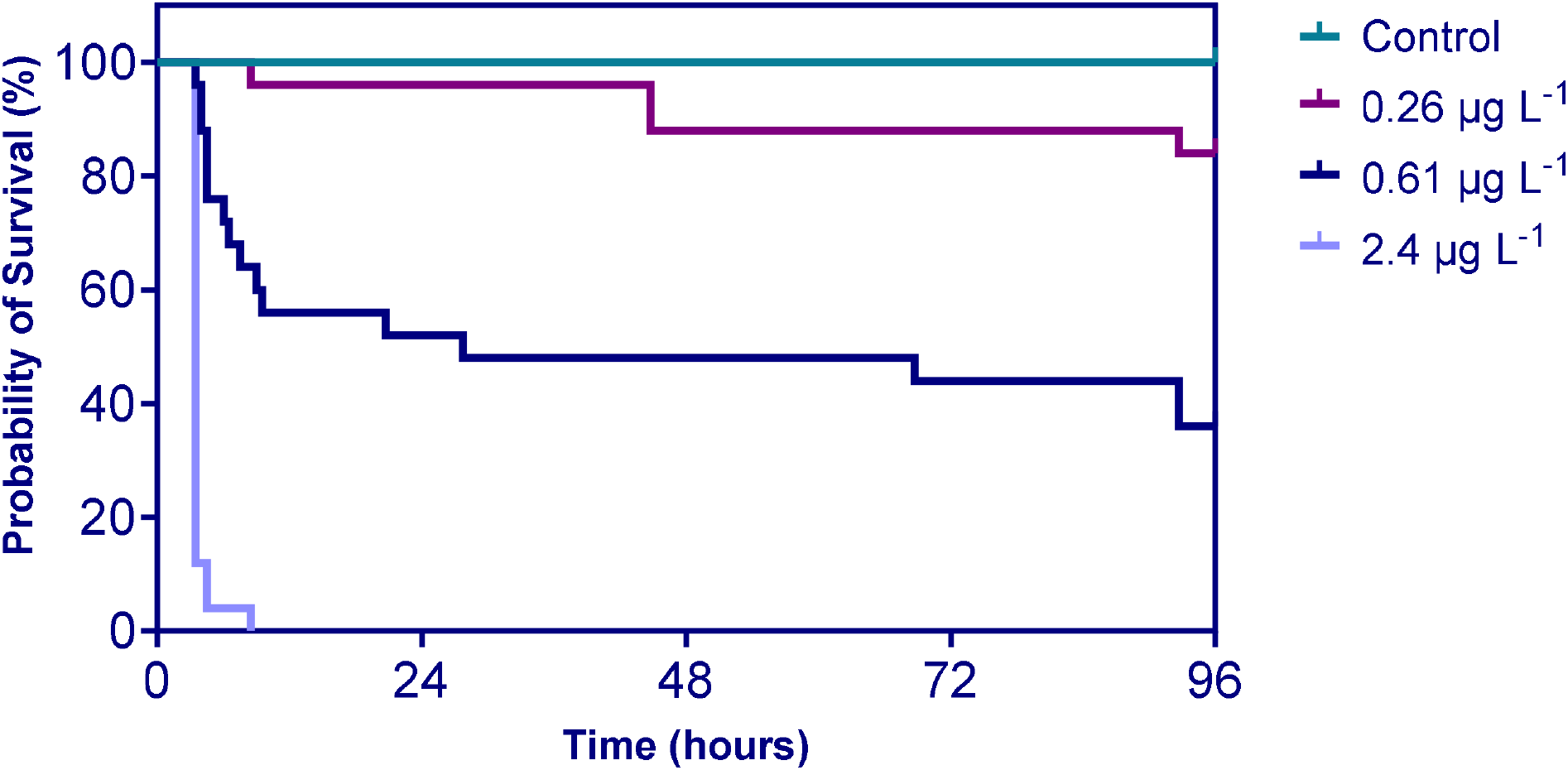
Survival time analysis for 6-week-old rainbow trout exposed to 6PPD-Q for 96 hours. Concentrations are time-weighted average measured concentrations.

**Figure 6.**
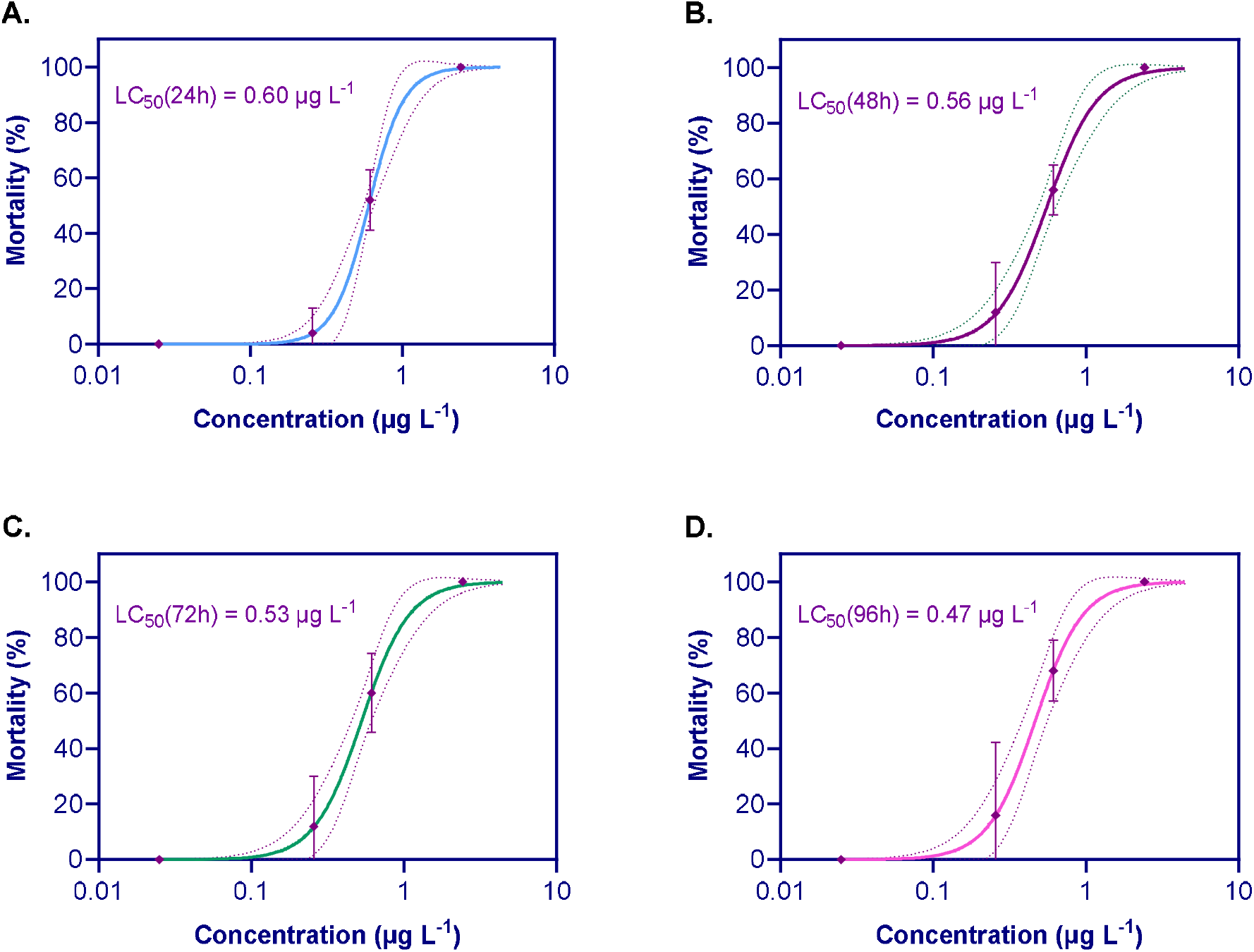
Percent mortality of 6-week-old rainbow trout exposed to 6PPD-Q at various times. The dotted lines indicate 95% confidence interval (CI). Points represent the mean of replicates for each concentration, standard deviation is represented by the bars. **A**. 24 hours (95% CI of 0.43 to 0.62 *μ*g/L); **B**. 48 hours (95% CI of 0.43 to 0.60 *μ*g/L); **C**. 72 hours (95% CI of 0.43 to 0.62 *μ*g/L); **D**. 96 hours (95% CI of 0.42 to 0.60 *μ*g/L).

This study has demonstrated that the early-life stages of rainbow trout are sensitive to 6PPD-Q and that 6 wph fry are the most sensitive life stage of this species studied thus far. This suggests that for those species in which adults exhibit sensitivity, there is a risk that the underdeveloped or earlier life stages may be even more sensitive. While the concern with URMS has largely centered around pre-spawn mortality, nursery sites where alevin and fry reside may also be subject to stormwater runoff, endangering the juveniles. This has far-reaching implications for population-level effects, as the loss of individuals at this stage of life greatly changes population dynamics (Milner et al., 2003).

### Conclusions and future outlook

Ultimately, this study found that post-swim-up rainbow trout fry are the most sensitive life stage of this species studied thus far, and that sub-chronic exposure to rainbow trout alevins results in mortality and developmental abnormalities that have not been observed in other life stages. The morphological deformities, as well as histopathologies, suggest that the gill is a key organ of toxicity. In addition, we observed an unusual abnormality in the caudal fin of alevins exposed sub-chronically, where blood pools throughout the fin. These sub-lethal effects suggest that, while mortality is key in determining benchmark concentrations, abnormalities which hinder a fish’s ability to grow and reproduce are also of concern to wild populations of species sensitive to 6PPD-Q. Sub-lethal effects must also be taken into account when characterizing the risk that 6PPD-Q poses to aquatic species.

While research has focused primarily on acute effects due to the pulsed nature of 6PPD-Q occurrence in the environment, more research is needed on the potential for chronic leaching of 6PPD-Q from TWP that were previously deposited within the sediment. Given that rainbow trout alevins are sensitive to 6PPD-Q and spend their early-life stage close to the sediment, there is a potential risk of these populations being exposed to continuous, low-dose 6PPD-Q exposure as TWPs settle into these environments. Detailed assessments of the potential leaching capacity of these settled particles are paramount to better characterizing and prioritizing sub-chronic and chronic risks versus those of pulsed exposure to contaminated roadway runoff.

As early-life stages of fishes are particularly sensitive to pollutants, this sensitivity allows for the detection of harmful effects at lower pollutant concentrations than might be observed in adult fishes. Thus, toxicity testing relying solely on adult fishes may not adequately protect the more sensitive early-life stages. Notably, in these experiments, post-swim-up fry exhibited the greatest response to 6PPD-Q, highlighting their vulnerability and the usefulness of this life stage in risk assessment. Swim-up fry exhibit similar behaviour to sub-adult fish, with the added benefits of decreased size and increased ease of husbandry, as well as potentially providing a more sensitive threshold for toxicity. The importance of including these life stages in risk assessments is key to ensuring comprehensive environmental protection of fish species from pollutants.

## Acknowledgements

This study was funded through a grant from Fisheries and Oceans Canada. Furthermore, the authors acknowledge support from the Discovery Grants of the Natural Sciences and Engineering Research Council of Canada (NSERC) to M.B. and M.H. M.B. is currently a faculty member of the Global Water Futures (GWF) program, which received funds from the Canada First Research Excellence Funds (CFREF). M.H. was supported by the Canada Research Chairs Program. The authors acknowledge the support of Zoë Henrikson, Dale Jefferson, and Azadeh Hatef (ATRF) in animal care. The authors also acknowledge the general lab support provided by Matthew Schultz, Julie Borsa, Kerstin Bluhm, Phillip Ankley, and Katherine Raes.

## Supplementary Material

**S1.**
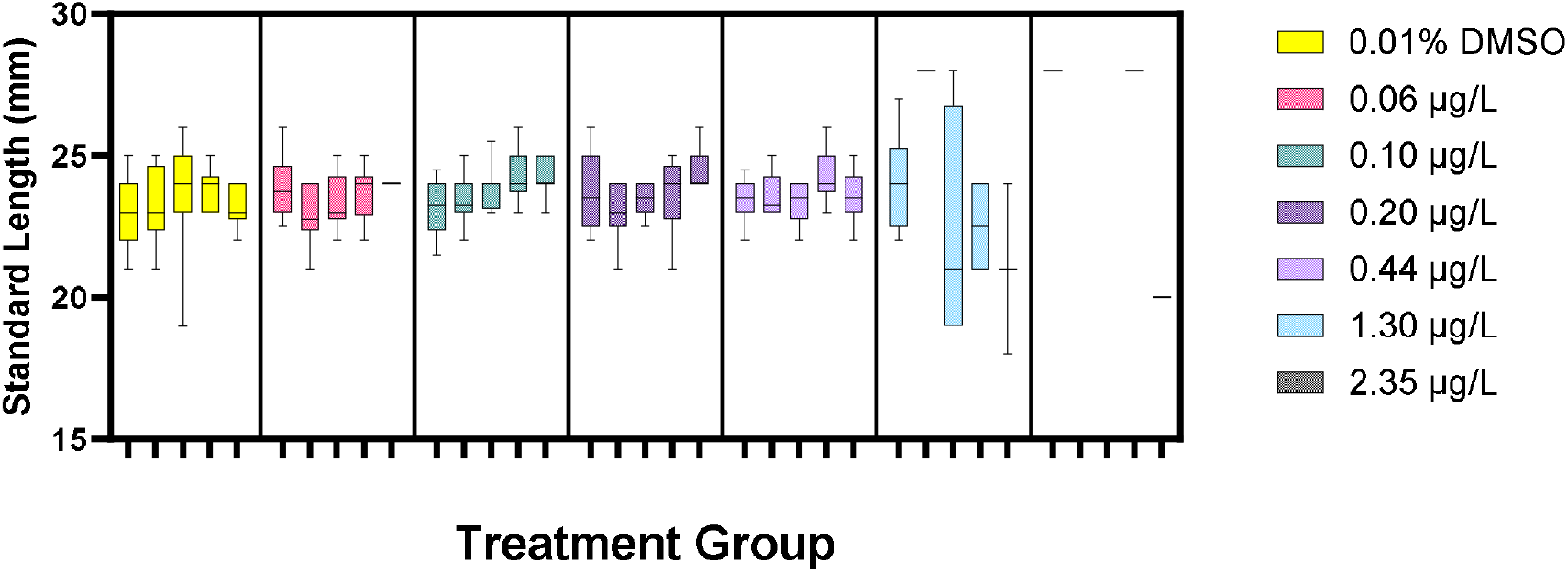
Average standard length (mm) per replicate of rainbow trout alevins exposed from hatch for 28 days to 6PPD-quinone

**S2.**
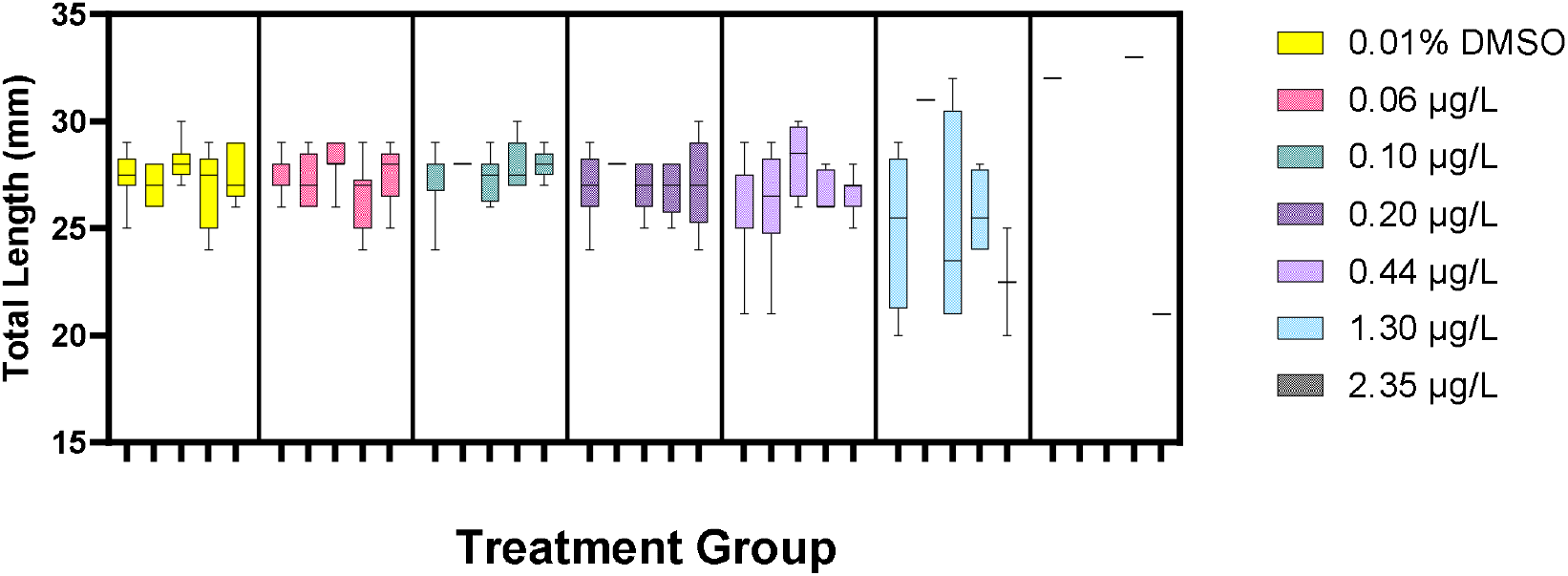
Average total length (mm) per replicate of rainbow trout alevins exposed from hatch for 28 days to 6PPD-quinone

**S3.**
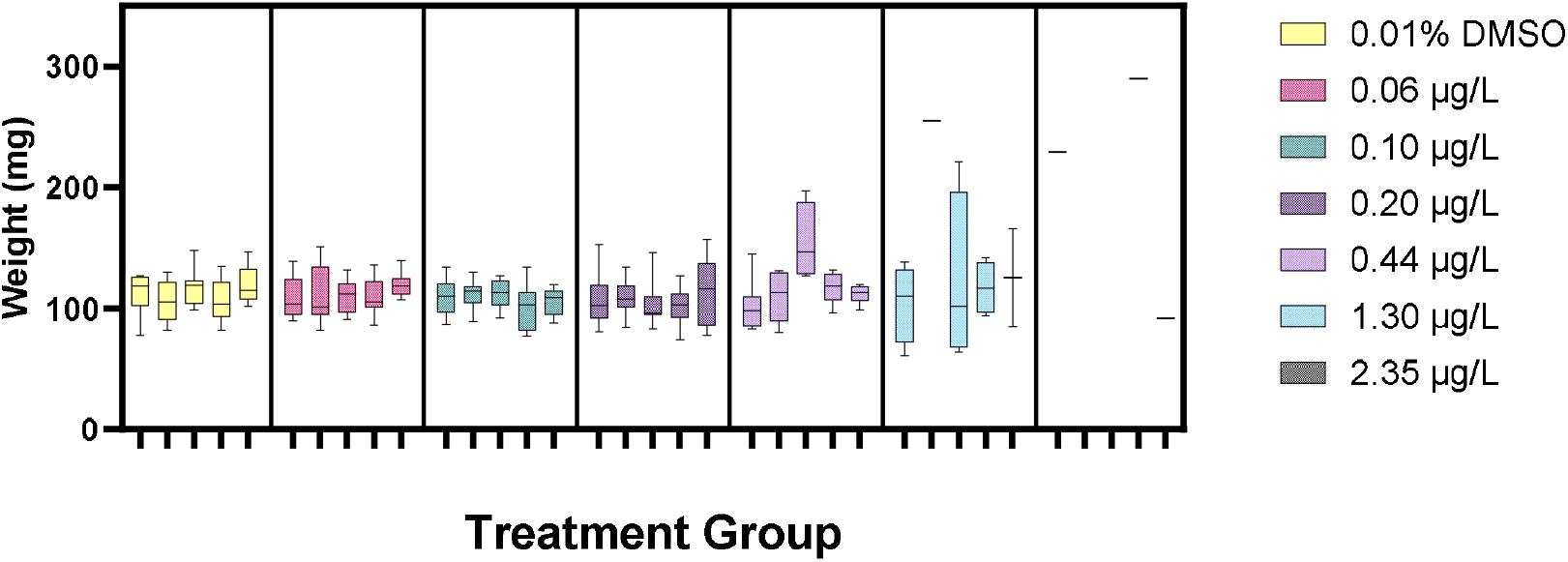
Average weight (mg) per replicate of rainbow trout alevins exposed from hatch for 28 days to 6PPD-quinone

**S4.**
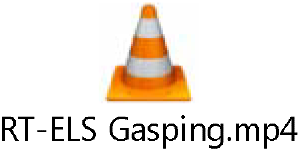
Example of behaviour consistent with URMS, observed in the juvenile acute study, including loss of coordination and gasping.

